# How Mitochondria Distribute and Align in HeLa Cells: A Computational Analysis On Electron Microscopy Images

**DOI:** 10.64898/2026.04.16.718887

**Authors:** Daniel Brito-Pacheco, Panos Giannopoulos, Constantino Carlos Reyes-Aldasoro

**Affiliations:** School of Science and Technology, City St. George’s, University of London, EC1V 0HB, London, United Kingdom; The Institute of Cancer Research, Integrated Pathology Unit, Division of Molecular Pathology, Sutton, United Kingdom

## Abstract

This paper investigates the way in which mitochondria distribute and align inside HeLa cells observed with serial block-face scanning electron microscopy. Four models of alignment were considered: (1) mitochondria exhibiting no discernible alignment pattern, (2) mitochondria aligned pointing towards the nucleus of the cell, (3) mitochondria aligned all in one direction when viewed from above, (4) mitochondria aligned tangent to the surface of the nucleus. These models were named (1) *unaligned*, (2) *petals*, (3) *racecars*, and (4) *clouds*. The mitochondria, nucleus and plasma membrane of 25 individual cells were segmented. A total of 12,299 mitochondria were identified and analysed. Alignment of the major axis of each mitochondrion was calculated in two ways: relative to a ray that joins it to the centroid of the nucleus, and relative to a ray that joins it to the nucleus’ surface. Results indicate that mitochondria tend to align tangentially to the nucleus surface, i.e., a *clouds* model. In addition, differences in the spatial distributions of the mitochondria were found and quantified with clearly defined metrics. The methodology here presented can be extended to other acquisition settings where the distribution and alignment of cells could be important, for instance, histopathology.

## 1 Introduction

Mitochondria are cellular organelles present in most eukaryotic cells and provide power to the cells through bioenergetic mechanisms [16]. Since their discovery in the late 1800’s, it has been understood that mitochondria play a fundamental role in cellular metabolism, energy production, and apoptosis [1, 3]. The spatial organisation of the mitochondria within the cytoplasm can influence physiological processes [12]. Thus, understanding how mitochondria distribute and align relative to the nucleus may therefore reveal new insights into the broader organisation of organelles inside a cell and how this affects the metabolism.

Advances in electron microscopy (EM) allow the observation of mitochondria and other subcellular structures with a much higher level of detail when compared to traditional microscopy. The images produced allow to detect, not only if mitochondria are present or not, but the exact position and orientation of the mitochondria with respect to the rest of the cell. Therefore, it is of interest to develop a methodology to identify individual mitochondria within cells and extract the geometric characteristics of their distribution and alignment.

In this work, a methodology to find the principal orientations and spatial distribution of mitochondria inside HeLa cells is presented. The methodology relies in previous works that provide accurate segmentation of nuclei, cellular membrane and mitochondria [2, 10]. Using volumetric EM data, the hypothesis that mitochondria exhibit a preferential alignment direction relative to the nucleus was studied. Four possible alignment models were conjectured:

– *Unaligned:* mitochondria do not present a discernible pattern of alignment, neither intrinsically nor extrinsically (Fig. 1 (a)).
– *Racecars:* mitochondria exhibit a preferential direction extrinsic to the cell. In simpler terms, from an outside perspective, the principal axes of mitochondria point in a parallel direction (Fig. 1 (b)).
– *Petals:* mitochondria align perpendicular to the nucleus’ surface, with their principal axes pointing toward the nucleus (Fig. 1 (c)).
– *Clouds:* mitochondria align tangent to the nucleus surface, with their principal axes pointing around the nucleus (Fig. 1 (d)).

**Fig. 1.**
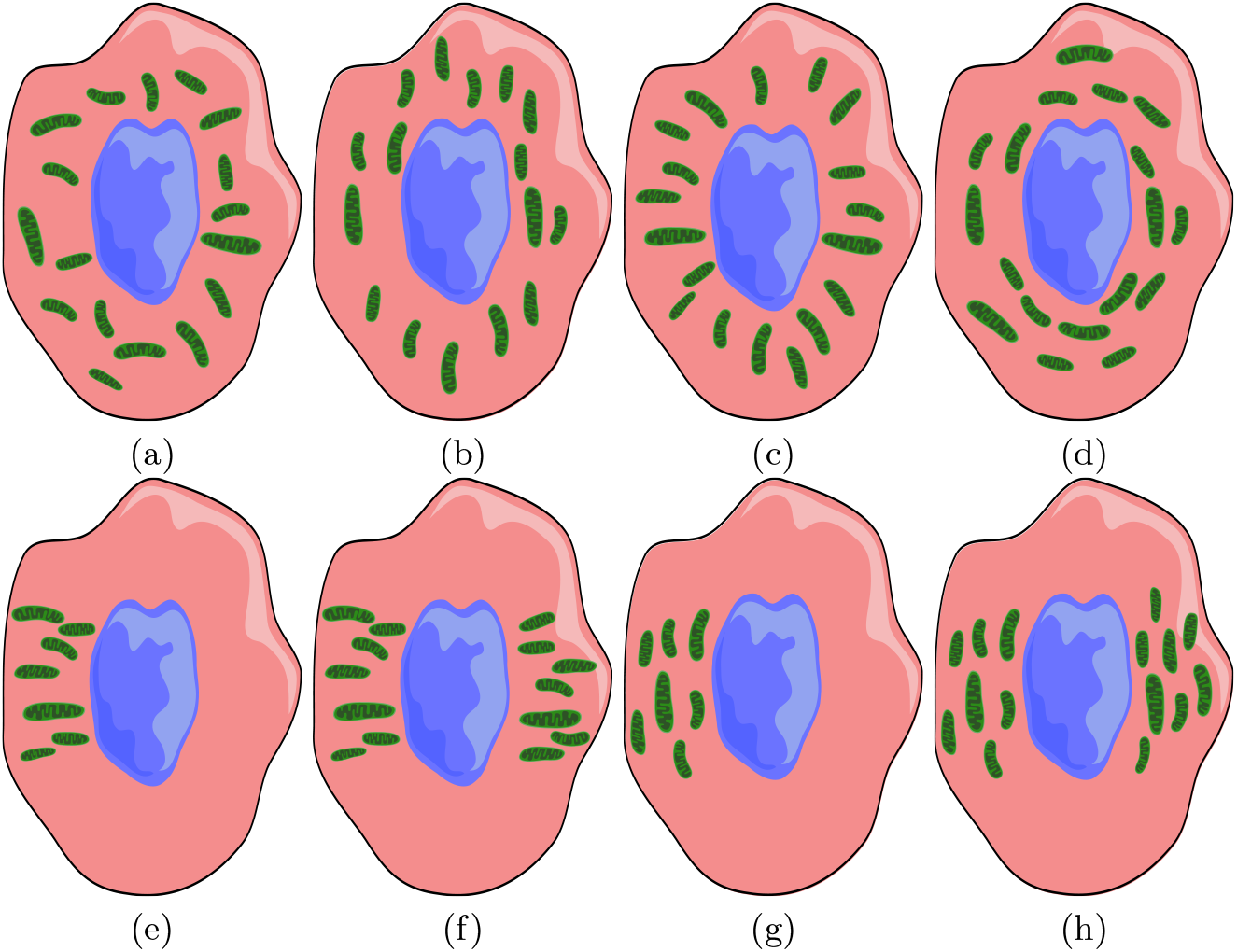
Four possible alignment models of the mitochondria inside a cell. (a) *Unaligned* model: mitochondria do not present a discernible pattern of alignment. (b) *Racecars* model: mitochondria aligned in one single direction when viewed from outside. (c) *Petals* model: mitochondria aligned pointing toward the nucleus’ surface. (d) *Clouds* model: mitochondria aligned tangent to the nucleus’ surface. It should be noted that when the mitochondria are not uniformly distributed around the cell, two models may be compatible. For example, in (e),(f) the *petals* model is not mutually exclusive with the *racecars* model: the principal direction of most mitochondria can point towards the nucleus, and, when distributed along a single axis, also show an extrinsic parallel alignment. (g),(h) Similarly, the *clouds* model is not mutually exclusive with the *racecars* model: the principal direction of most mitochondria can be parallel to the nucleus’ surface, and, also show an extrinsic parallel alignment.

It should be considered that some of these models are not mutually exclusive, i.e., the *racecars* model, and either *petals* or *clouds* could be satisfied depending on the distribution of the mitochondria. The mitochondria of a cell can exhibit an intrinsic perpendicular-to-the-surface or parallel-to-the-surface alignment and simultaneously show an extrinsic parallel alignment (Fig. 1 (e)-(h)).

The spatial distribution of the mitochondria around the nucleus was also quantified computationally, thereby contributing to a better understanding of intracellular organisation.

## 2 Materials and Methods

### 2.1 Materials

The preparation of the cells and the basic segmentation steps have been published earlier. Briefly, HeLa cells were prepared following the protocol detailed in [15]. The cells were embedded in Durcupan resin and images were captured using a serial block-face scanning electron microscope as described in [7]. The dataset consists of a volume of 8000×8000×517 voxels (Fig. 2 (a)) and is openly available via EMPIAR (https://www.ebi.ac.uk/empiar/EMPIAR-10094/). Given the dataset resolution of 10nm×10nm×50nm, the volume was rescaled to isotropic units for analysis.

**Fig. 2.**
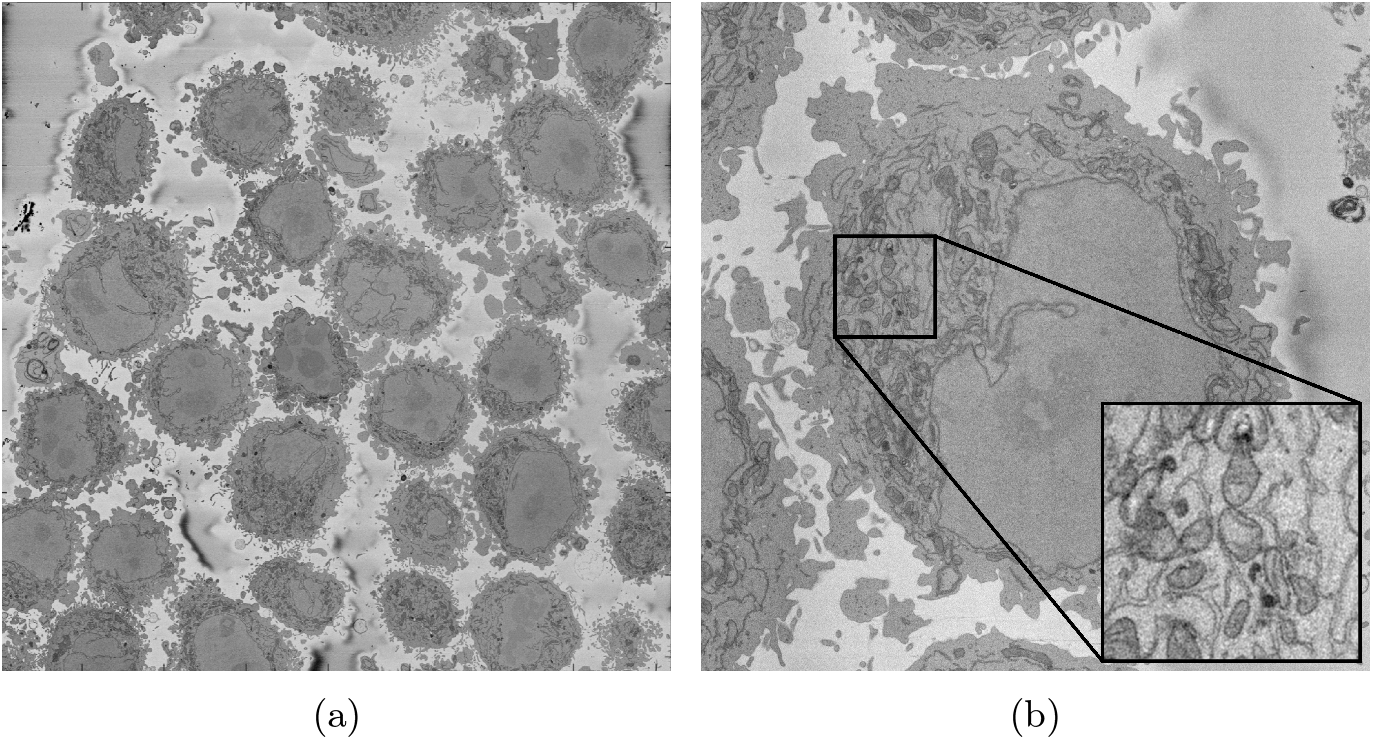
(a) A single slice of the original 8000×8000×517 volume showing several HeLa cells. (b) A slice of a region-of-interest centred on a single HeLa cell and zoom in on a region of the slice containing mitochondria.

### 2.2 Methods

#### Segmentation of HeLa Cells

Regions-of-Interest (ROIs) were automatically detected for 25 cells in the volume (Fig. 2 (b)) and nuclei, plasma membranes and mitochondria automatically segmented (Fig. 3). For each cell, the nucleus and plasma membrane were segmented via the processes described in [10] and [11], respectively. The mitochondria were segmented using the MitoNet Deep Learning model, accessed through a Python package called *empanada* [6]. Four segmentation models were tested and compared [2]. Of these methods, MitoNet obtained the highest score (0.659) when tested on a 2-dimensional benchmark dataset. Although the theoretical maximum of the Jaccard index is 1, an inter-observer (human-to-human) comparison produced a score of 0.696. This value provides a more realistic upper bound for performance on this task. Relative to this, the score obtained by MitoNet corresponds to 94.68% of the inter-observer agreement. This indicates that the model performs close to the level of human annotators.

**Fig. 3.**
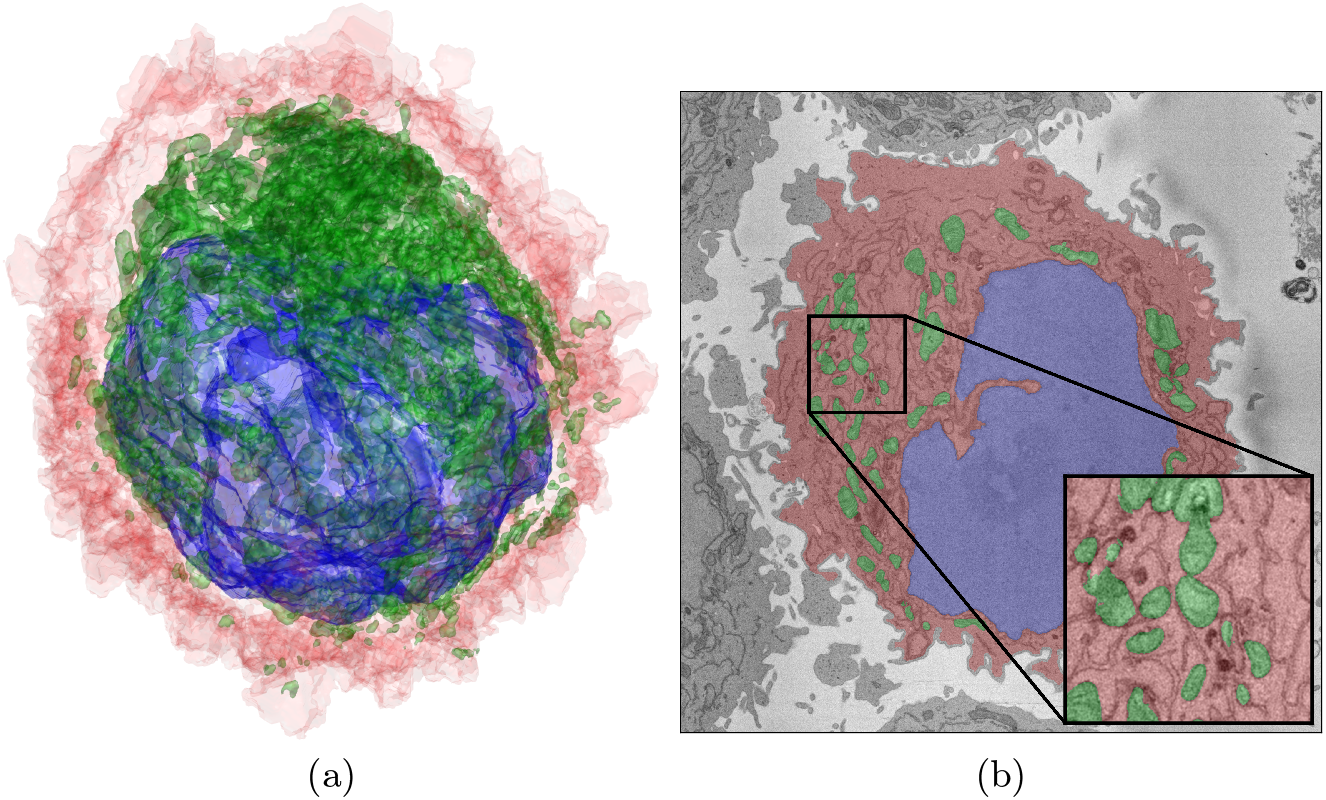
Segmentation of the cellular structures. For each cell, the following structures were segmented: nucleus (blue), cell (red, region contained within the plasma membrane), mitochondria (green). Whilst there is a single instance of nucleus and cell, each mitochondria is considered as a separate instance.

**Fig. 4.**
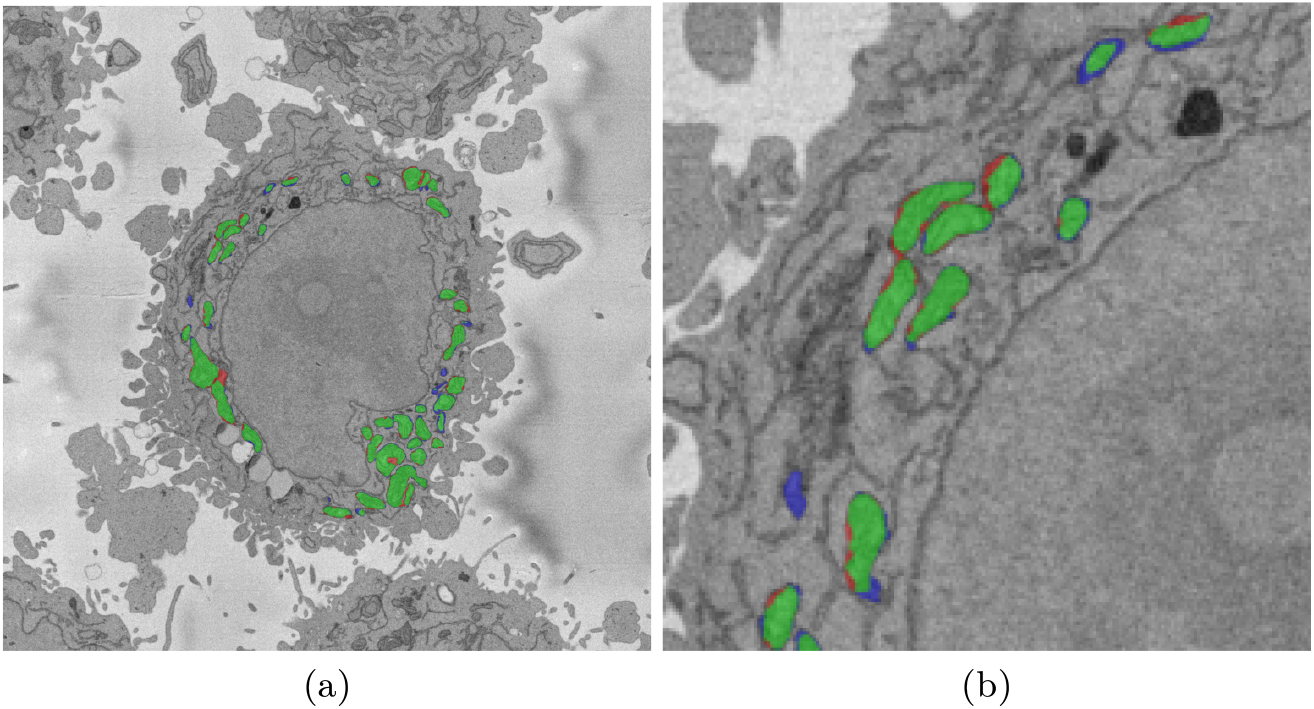
Illustration of the mitochondria segmentation accuracy (a) one slice of a HeLa cell, (b) region of interest. For both cases green pixels indicate True Positive (TP), red pixels False Positives (FP), and blue pixels False Negatives (FN). It can be seen that errors occur around the edges of the mitochondria and in some cases (bottom left of (b)) a whole mitochondrion was missed. For the purposes of this work, i.e., to analyse the orientation and alignment of the mitochondria, the orientation of the each mitochondrion will not be heavily influenced by the errors detected.

#### Directional Statistics

Before proceeding to following subsections, it is necessary to outline the basic concepts behind directional statistics. Statistics on directional measurements (angles) differ from traditional statistics since, for most applications, angles are measured using mod 360° (2*π* radians). The angles *θ*_1_ = 10° and *θ*_2_ = 350° point in very similar directions. Intuitively, their mean is expected to be 0° = 360° (the angle halfway between them), however a traditional computation yields 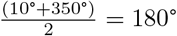.

To get a better representation of a mean, one must adopt the language of complex numbers. In this context, a unitary vector in 2D Euclidean Space with angle *θ* (measured anti-clockwise from the *x*-axis) is given as the complex number *z* = *e*^*iθ*^ = cos *θ* + *i* sin *θ*. Thus, the sample {*θ*_1_, …, *θ*_*N*_} of *N* angles, is equivalent to a sample of complex numbers 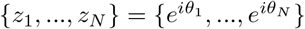. The *n*^th^ moment of the sample is defined as:

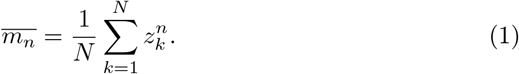

Then, the resultant vector of a sample of *N* unitary vectors is the first moment of the sample:

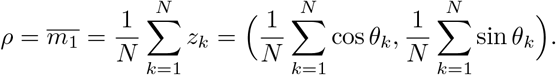

The circular mean angle is the angle formed by *ρ* and the *x*-axis, often given by the arctan2 function […]:

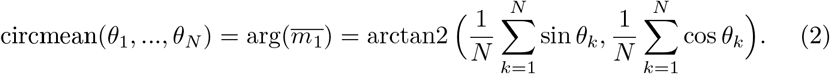

The length of the resultant vector is

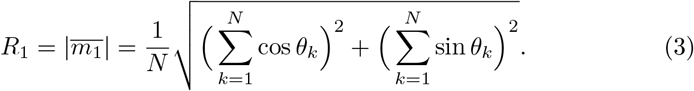

Analogously, the length of the second moment is given by:

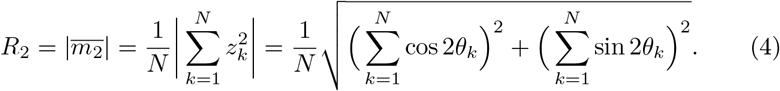

Finally, a measure of dispersion is the circular standard deviation *σ*. It must be noted that, unlike the linear standard deviation for a normal distribution, it does not admit a simple probabilistic interpretation (such as the fact that approximately 68% of observations lie within one standard deviation of the mean). Instead, *σ*, defined below reflects variability through the mean resultant length *R*_1_:

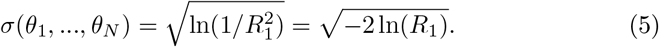

As *R*_1_ approaches 1, *σ* approaches 0, reflecting low variability in the sample; conversely, as *R*_1_ tends to 0, *σ* diverges to +∞, indicating high variability. Since directional data are periodic, defined modulo 2*π*, this measure does not allow for a straightforward notion of “distance in standard deviations” from the mean.

#### Orientation of Mitochondria

Each cell *C*_*k*_ contains *M*_*k*_ mitochondria. In this subsection, the values that were measured to determine the model which best represents the alignment of mitochondria inside a cell are described:

– *α*_*ki*_: the angle formed by the mitochondrion’s principal axis *a*_*ki*_ and the line which joins its centroid *m*_*ki*_ to its nucleus centroid (centroid-to-centroid line).
– *β*_*ki*_: the angle formed by the mitochondrion’s principal axis *a*_*ki*_ and the line which joins its centroid *m*_*ki*_ to the closest point on its nucleus surface *p*_*ki*_.
– *γ*_*xki*_, *γ*_*yki*_, *γ*_*zki*_ : the angles formed by the mitochondrion’s principal axis *a*_*ki*_ and the coordinate *x, y, z* axes, respectively.

The values *α* and *β* are exclusively intrinsic to the cell’s nucleus and mitochondria; and are intended to discriminate between the *clouds* and *petals* models. As mentioned in Section 1, the *petals/clouds* and *racecars* models are not mutually exclusive; thus, *γ*_*x*_, *γ*_*y*_, *γ*_*z*_, which are values extrinsic to the cell (mitochondrial directions are compared against global axes) reveal whether there exists a preferential alignment of mitochondria along fixed spatial directions. In particular, the standard deviations of these values in a given cell reveal the variability of mitochondrial alignment relative to the external coordinate system.

The angle, *α*_*ki*_, was computed as the angle between *a*_*ki*_ and the vector *m*_*ki*_−*n*_*k*_. Similarly, was computed as the angle formed by *a*_*ki*_ and the vector *m*_*ki*_−*p*_*ki*_. It must be noted that *α* and *β* take values between 0 and 180°. In this context, 0° and 180° point in the same direction, so to compute the mean direction, the means mod 180° of the double angles were computed from the definition in (2):

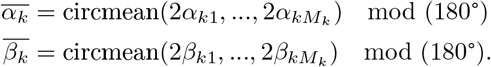

Rayleigh tests of uniformity are statistical tests that evaluate the null hypothesis, *H*_0_, that a population of angles is uniformly distributed around the circle, against the alternative hypothesis, *H*_1_, that the angles exhibit a preferred direction, and are therefore not uniformly distributed […]. This transformation maps the data onto the full circular domain [0, 360°), allowing the test to more faithfully detect preferential alignment.

Additionally, the circular means of the average angles across cells were computed as:

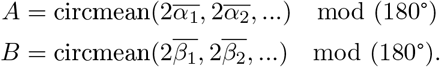

To allow for generalisation to the bigger population of HeLa cells, confidence intervals at 95% for *A* and *B* were estimated by bootstrapping the measured 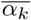 and 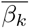 across all 25 cells, with 5000 bootstrap samples. Similar to *α* and *β, γ*_*x*_, *γ*_*y*_, *γ*_*z*_ take values between 0° and 180°. Their dispersion is therefore assessed using doubled angles in the formula in (5) then divided by two.

All computations were done in Python, directly from the volumes of segmented isotropic voxels. As well as the principal axis *a*_*ki*_, the intermediate *b*_*ki*_ and minor axes *c*_*ki*_ were also computed. All three were computed using Principal Component Analysis (PCA). Mitochondria with an aspect ratio (|*a*_*ki*_|*/*|*c*_*ki*_|) less than two were not included as near-spherical mitochondria may introduce noise to the computations.

#### Spatial Distribution of Mitochondria

To study the position of mitochondrial centroids *m*_*ki*_ relative to the nucleus, the origin of the 3D Euclidean space was centred at the centroid of the nucleus. Three quantities were measured: the polar azimuth coordinates obtained by projecting *m*_*ki*_ to each of the *xy, xz, yz* coordinate planes. These coordinates are denoted by 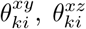, and 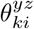, respectively. *R*_1_ and *R*_2_ were then computed from the resulting sets of azimuthal measurements.The notation 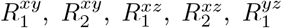, and 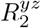 indicates the corresponding projection plane.

In this context, *R*_1_ captures unimodal concentration in location around the nucleus. As the length of the resultant vector, it approaches 1 when all mitochondria are close together, and approaches 0 when mitochondria are distributed uniformly or symmetrically. *R*_2_ captures biaxial symmetry; it approaches 1 if mitochondria are distributed in two diametrically opposite groups.

It is important to note that neither *R*_1_ nor *R*_2_ can single-handedly capture the distribution of mitochondria around the nucleus; rather, both must be considered to characterise the dispersion relative to the nuclear centroid. Consider a toy example where two mitochondria, *m*_1_ and *m*_2_ are placed at azimuths of 0° and 80°. This distribution is clearly one-sided, the values of *R*_1_ = 0.77, *R*_2_ = 0.17, reflect this. However, a tighter cluster, where 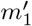 and 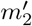 are located at 15° and 30°, respectively, yields a much higher *R*_2_ = 0.87, which could misleadingly suggest biaxial symmetry. By also considering *R*_1_ = 0.97 the picture becomes clearer. The high value of *R*_1_ outweighs the apparent bisymmetry suggested by *R*_2_. In practice it is advisable to first examine *R*_1_ to assess unimodality. Only when *R*_1_ is small, should one turn to *R*_2_ to differentiate between uniform and bisymmetrical distributions.

## 3 Results

### 3.1 Orientation

The normalised frequencies of the orientation of the 12,299 mitochondria are presented as rose plots for *α*_*i*_ and *β*_*i*_ (Fig. 5 (a), (b)). The average angles 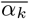, 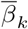 are shown in Table 1. All Rayleigh tests yielded *p*-values which were much smaller than 0.0001. The across-cell results yielded *A* = 89.74°, *B* = 88.90° with 95% confidence intervals of [87.11°, 92.16°] and [86.16°, 91.27°], respectively.

**Table 1.** All computed values for mitochondria in their respective cells. 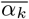: mean angle formed by the principal axis and the centroid-to-centroid line. *s*(*α*): standard deviation of angles formed by the principal axis and the centroid-to-centroid line. 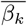:mean angle formed by the principal axis and the centroid-to-surface line. *s*(*β*): standard deviation of angles formed by the principal axis and the centroid-to-surface line. max *s*(*γ*): maximum standard deviation of angles formed by the principal axis and the coordinate axes. 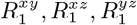: length of the mean of the azimuths when mitochondria centroids are projected to the *xy, xz, yz* planes, respectively (See Section 2.2). 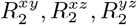: length of the mean of the double azimuths when mitochondria are projected to the *xy, xz, yz* planes, respectively (See Section 2.2).

| Cell | $M_k$ | Orientation Statistics | | | | | Distribution Statistics | | | | | |
| --- | --- | --- | --- | --- | --- | --- | --- | --- | --- | --- | --- | --- |
| | | $\bar{\alpha}$ | $s(\alpha)$ | $\bar{\beta}$ | $s(\beta)$ | $\max s(\gamma)$ | $R_1^{xy}$ | $R_1^{xz}$ | $R_1^{yz}$ | $R_2^{xy}$ | $R_2^{xz}$ | $R_2^{yz}$ |
| 01 | 314 | 89.89° | 0.94 | 89.80° | 0.95 | 1.54 | 0.41 | 0.35 | 0.28 | 0.08 | 0.27 | 0.14 |
| 02 | 329 | 90.18° | 0.95 | 89.88° | 0.89 | 1.49 | 0.11 | 0.09 | 0.10 | 0.19 | 0.27 | 0.21 |
| 03 | 549 | 92.74° | 1.05 | 93.37° | 0.95 | 1.38 | 0.79 | 0.77 | 0.27 | 0.46 | 0.43 | 0.04 |
| 04 | 347 | 91.39° | 1.00 | 90.26° | 1.01 | 1.61 | 0.08 | 0.18 | 0.12 | 0.49 | 0.15 | 0.46 |
| 05 | 328 | 83.55° | 1.08 | 88.25° | 0.95 | 1.46 | 0.36 | 0.40 | 0.41 | 0.22 | 0.32 | 0.27 |
| 07 | 338 | 88.79° | 0.88 | 85.44° | 0.91 | 1.37 | 0.24 | 0.34 | 0.29 | 0.27 | 0.09 | 0.04 |
| 08 | 635 | 91.25° | 1.07 | 91.33° | 1.03 | 1.64 | 0.36 | 0.04 | 0.34 | 0.10 | 0.21 | 0.13 |
| 09 | 621 | 98.26° | 1.06 | 95.75° | 1.04 | 1.56 | 0.19 | 0.30 | 0.34 | 0.23 | 0.12 | 0.09 |
| 10 | 449 | 88.29° | 0.94 | 85.63° | 0.91 | 1.65 | 0.53 | 0.44 | 0.45 | 0.23 | 0.12 | 0.24 |
| 11 | 311 | 99.15° | 1.15 | 94.79° | 0.90 | 1.29 | 0.12 | 0.26 | 0.25 | 0.06 | 0.41 | 0.33 |
| 12 | 598 | 94.82° | 1.25 | 93.97° | 1.14 | 1.49 | 0.63 | 0.57 | 0.50 | 0.38 | 0.21 | 0.13 |
| 13 | 606 | 96.02° | 1.06 | 93.01° | 0.96 | 1.29 | 0.16 | 0.18 | 0.10 | 0.13 | 0.10 | 0.23 |
| 16 | 428 | 81.97° | 1.03 | 83.90° | 0.93 | 1.45 | 0.30 | 0.39 | 0.30 | 0.06 | 0.18 | 0.26 |
| 17 | 609 | 100.03° | 1.00 | 97.49° | 0.91 | 1.31 | 0.47 | 0.33 | 0.36 | 0.25 | 0.21 | 0.16 |
| 18 | 604 | 78.85° | 1.09 | 78.69° | 1.02 | 1.46 | 0.48 | 0.39 | 0.56 | 0.25 | 0.17 | 0.18 |
| 19 | 554 | 95.36° | 1.14 | 96.10° | 1.03 | 1.73 | 0.34 | 0.38 | 0.32 | 0.05 | 0.28 | 0.13 |
| 20 | 563 | 73.00° | 1.01 | 69.67° | 0.97 | 1.38 | 0.16 | 0.70 | 0.66 | 0.07 | 0.40 | 0.30 |
| 21 | 442 | 87.02° | 1.02 | 82.99° | 0.88 | 1.43 | 0.05 | 0.26 | 0.25 | 0.21 | 0.09 | 0.10 |
| 22 | 394 | 86.15° | 0.86 | 85.43° | 0.86 | 1.46 | 0.43 | 0.51 | 0.50 | 0.21 | 0.09 | 0.23 |
| 23 | 415 | 83.46° | 0.96 | 82.85° | 1.00 | 1.51 | 0.66 | 0.64 | 0.59 | 0.23 | 0.30 | 0.34 |
| 24 | 342 | 85.82° | 1.17 | 82.89° | 1.04 | 1.52 | 0.23 | 0.34 | 0.29 | 0.13 | 0.05 | 0.03 |
| 25 | 877 | 93.58° | 1.05 | 91.28° | 0.99 | 1.59 | 0.17 | 0.09 | 0.16 | 0.30 | 0.16 | 0.11 |
| 26 | 421 | 85.27° | 0.97 | 87.74° | 0.84 | 1.35 | 0.33 | 0.30 | 0.38 | 0.09 | 0.17 | 0.19 |
| 29 | 473 | 97.7° | 1.16 | 100.05° | 1.01 | 1.48 | 0.32 | 0.57 | 0.59 | 0.22 | 0.19 | 0.19 |
| 30 | 437 | 90.29° | 1.11 | 90.68° | 1.13 | 1.54 | 0.58 | 0.18 | 0.59 | 0.16 | 0.19 | 0.32 |

**Fig. 5.**
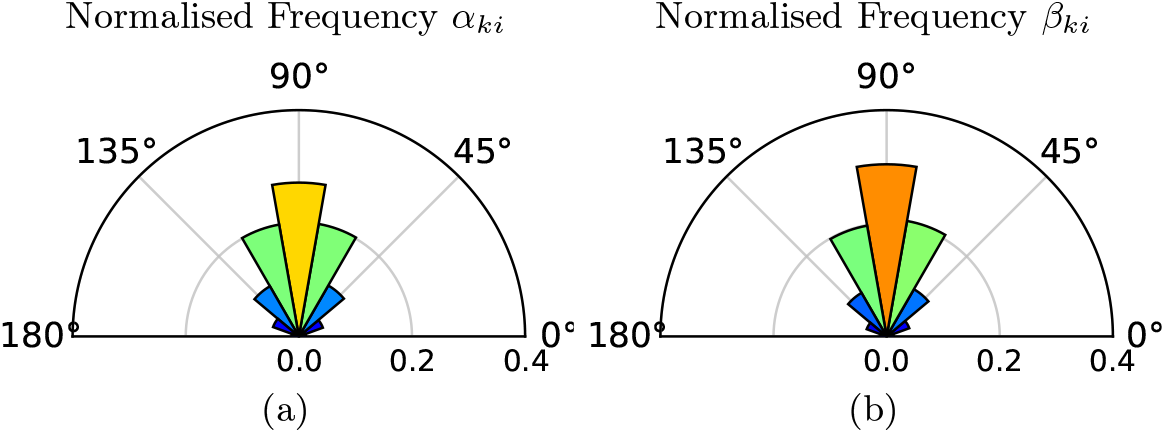
Normalised frequencies of (a) *α*_*ki*_ and (b) *β*_*ki*_ of all mitochondria across all 25 cells.

These diagrams and values clearly show that the majority of mitochondria align as the *cloud* model with angles close to 90° with respect to the rays from the centroid and with an even higher alignment when measured from the surface of the nuclei.

### 3.2 Distribution

Results for the computed values of 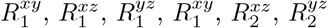 are shown in Table 1. Additionally, rose plots for 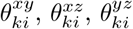 with corresponding *R*_1_ and *R*_2_ values for particular cases are highlighted in Fig. 7.

## 4 Discussion

The initial objective of this study was to identify how mitochondria are aligned and distributed relative to the nucleus inside HeLa cells, as seen with Electron Microscopy images using a clearly defined methodology. The computed values for *A* and *B*, with their 95% confidence intervals presented in 3.1 indicate that the major axes of mitochondria are predominantly perpendicular to the centroid-to-centroid and centroid-to-surface lines. These results point strongly towards a *clouds* model for the alignment of mitochondria inside HeLa cells. This is supported by Fig. 5 (a), (b) and Fig. 6 (a) where, the modes of *α*_*ki*_ and *β*_*ki*_ were close to 90°.

**Fig. 6.**
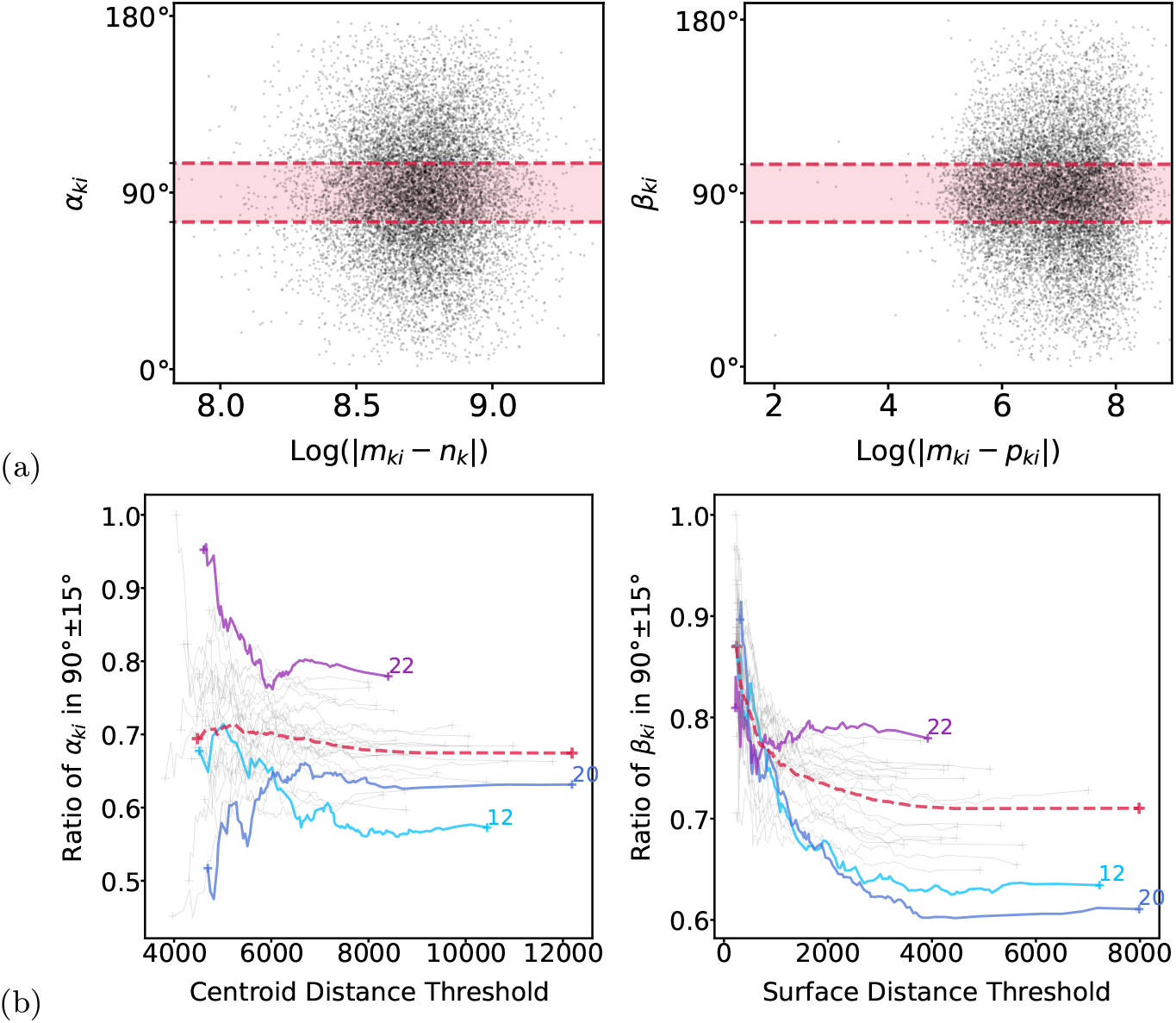
(a) Log of distance from mitochondria centroid to nucleus centroid (log(|*m*_*ki*_ − *n*_*k*_|)) vs. *α*_*ki*_ (left), and log of distance from mitochondria centroid to closest point on nucleus surface (log(|*m*_*ki*_, *p*_*ki*_|)) vs. *β*_*ki*_ (right). (b) Ratio of mitochondria within the angular interval 90° *±* 15° as a function of thresholds on the distance from the centroid (left) and the surface (right) of the nucleus for each cell (solid lines), and the complete volume (dotted line).

Moreover, it is clear that the orientations of mitochondria are more strongly affected by the surface of the nucleus, than the centroid (Fig. 5 (a) vs (b)). This suggests that as the surface of the nucleus varies from a perfect sphere, the mitochondria follow the variations.

The distance from the nucleus was also found to have an effect on the alignment of the mitochondria. As mitochondria get further away from the nucleus, the spread of both *α*_*ki*_ and *β*_*ki*_ increases (Fig. 6 (a)). The behaviour of the alignment of mitochondria based on distance in each cell (solid lines) and the entire volume (dotted line) was further captured in Fig. 6 (b). This finding correlates with the literature that has reported that mitochondria tend to be aligned along microtubules [14]. Thus, it could be inferred that the microtubules in the cytoplasm also tend to have an orientation around the nuclei of HeLa cells and can also have an influence on their shape [8]. For the entire volume, *α*_*ki*_ behaved relatively stable, the spread of values did not change as distance increased. More notable was the behaviour of *β*_*ki*_ with respect to the distance from the surface of the nucleus. For most cells, and on average, as mitochondria get farther away, the ratio that have a *β*_*ki*_ close to 90° becomes lower. However, for individual cells this was not always the case: three cells were highlighted (*12, 20, 22*). Cells *12* and *20* reached the lowest ratio of mitochondria within 15°of 90°for *α*_*ki*_ and *β*_*ki*_, respectively. For values of *α*_*ki*_, the curve of cell *12* follows a similar pattern to the global curve, whereas cell *20* shows an uncommon behaviour, starting low, then having a higher ratio closer to 90°. In contrast, cell *22*, had the highest ratio of mitochondria close to 90° for both *α*_*ki*_ and *β*_*ki*_. Additionally, it seemed to have a spike of perpendicularly aligned mitochondria after some distance from the nucleus. These differences may be related with the observed correlation of mitochondrial location and structure and its dependence on environmental factors like CO2 availability [9] or cell cycle stage [13].

The distribution of mitochondria around the nucleus was also described, *R*_1_ and *R*_2_ captured the differences in distributions. Cells where the distribution was one-sided, when viewed from above, yielded high values for *R*_1_ (Fig. 7 (a), (d)). Low values for *R*_1_ were obtained for cells with more uniform or symmetric distributions (Fig. 7 (b), (c)). *R*_2_ yielded high values for two-sided mitochondrial distributions (Fig. 7 (b)), and both values were low when the distribution was close to being uniform (Fig. 7 (c)). It is worth noting that even though comparable values for *R*_2_ can be obtained for one-sided and two-sided distributions, as is the case for Fig. 7 (a), (b), the combination of both metrics is what reveals the differences in spatial distribution, seen in the very small *R*_1_ value for Fig. 7 (b). Again, this correlates with the literature that has reported that mitochondria, especially during the interphase in HeLa cells, are not distributed uniformly around the cell, but tend to aggregate more densely around the nucleus (peri-nuclear region) than the cell periphery [5].

**Fig. 7.**
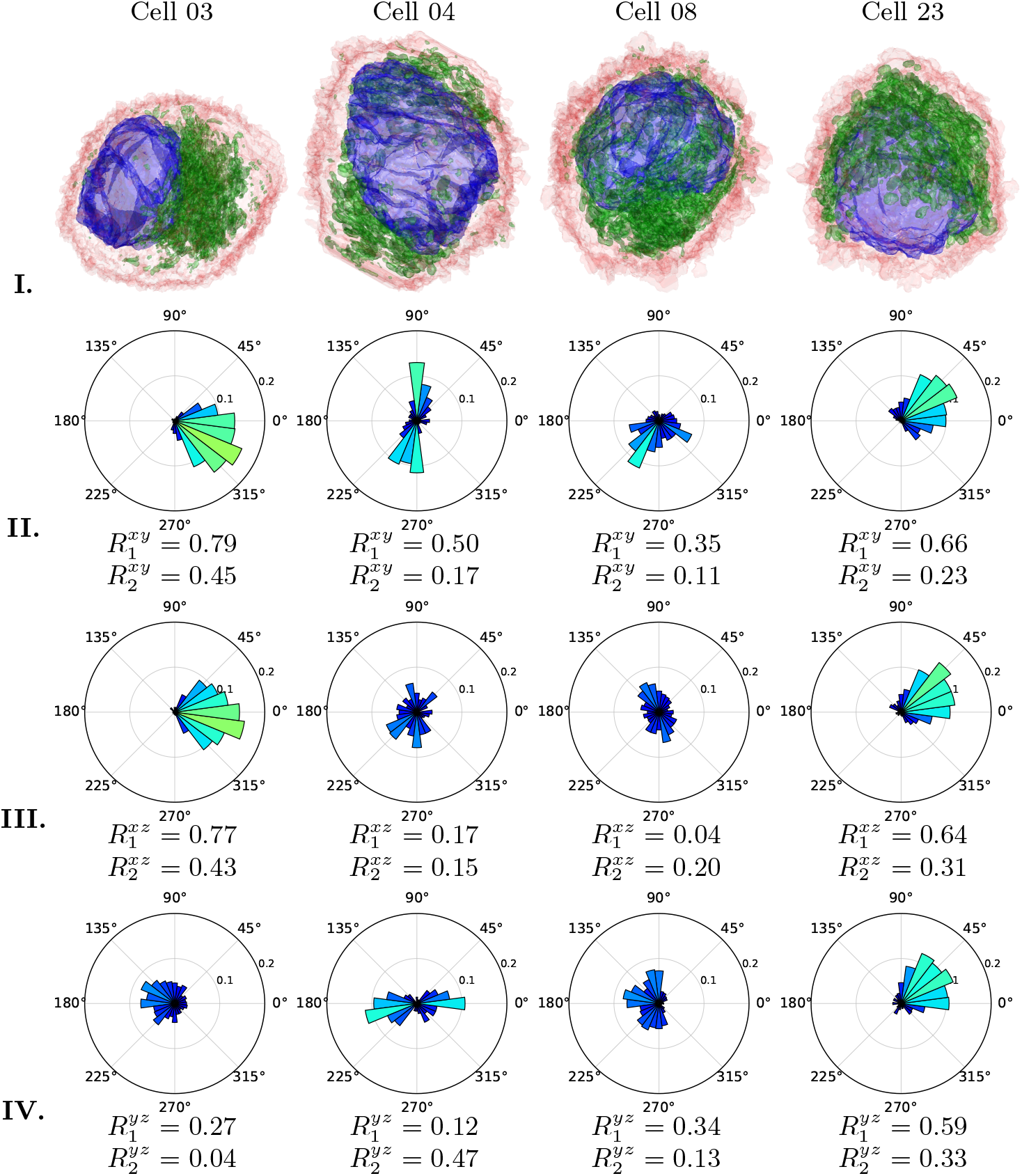
**I**. Segmentations of mitochondria, nucleus and membrane for four distinct cells (03, 04, 08, 23). **II.-IV**. Rose plots and computed values for *R*_1_, *R*_2_ capture the distributions of mitochondria around the centroid of the nucleus. Projecting on all three coordinate planes to compute *R*_1_ and *R*_2_ values reveal differences in one and two-sidedness of the mitochondria clouds.

## 5 Conclusion

In this work, a method to describe the orientations and spatial distribution of mitochondria inside HeLa cells was presented. The results strongly pointed towards a model where mitochondria align their major axis tangent to the nucleus’ surface (*cloud* model). Qualitatively, it was also observed that *R*_1_ and *R*_2_ capture differences on the spatial distributions of mitochondria around the nucleus.

The qualitative evaluations performed in this study show that the method is robust and is a step towards understanding the organisation of organelles inside cells. The main contribution presented here was that of quantifying mitochondrial alignment and distribution precisely, through the use of EM images and accurate computational methods.

## Future Work

This work established a computational method for quantifying mitochondrial organisation within cells. Future efforts will focus on two things: (1) expanding the analysis beyond normal HeLa cells to include different treatments or pathological conditions and (2) building on the mathematical insight to capture more complex geometries and subcellular organisation. For the latter, recent advances in describing the structure of mitochondrial organisation through topological data analysis have been made [4]. The combination of sophisticated topological descriptors with different treatments of cells, could uncover how disease affects the internal organisation of cells.

## Code and Data Availability

Code and data are freely available. HeLa images are available from *http://dx.doi.org/10.6019/EMPIAR-10094*, code from *https://github.com/reyesaldasoro/MitoEM*, and segmented nuclei, cells, and mitochondria from *https://github.com/reyesaldasoro/HeLa_Cell_Data*.

